# Establishment of Integration-Free iPSCs from Diverse Porcine Species: A Novel Resource for Conservation and African Swine Fever Research

**DOI:** 10.64898/2026.05.02.722394

**Authors:** Qiuye Bao, Christina Yingyan Lim, Nicole Liling Tay, Hui Li Yeo, Kanchana Punyawai, Chia-da Hsu, Shin Min Chong, Shangzhe Xie, Jonathan Yuin-Han Loh, Soon Chye Ng, Oz Pomp

## Abstract

The accelerating biodiversity crisis, compounded by emerging infectious diseases like African swine fever (ASF), necessitate innovative conservation and disease management. ASF susceptibility varies wildly across species, from near-100% mortality in Asian suids to asymptomatic carriage in African forest species. We report the first successful derivation of integration-free induced pluripotent stem cells (iPSCs) from four phylogenetically distinct species: wild boar (*Sus scrofa*), Bornean bearded pig (*Sus barbatus*), Babirusa (*Babyrousa babyrussa*), and Red river hog (*Potamochoerus porcus*). Using Sendai virus-mediated reprogramming, we achieved efficiencies between 0.003% and 0.26%. These iPSCs were successfully differentiated into CD14⁺CD11b⁺ monocytes - the primary target cells for the ASF virus - establishing a renewable, comparative research platform. This system enables host-pathogen studies previously hindered by ethical and logistical constraints of wildlife sampling. Beyond disease research, these iPSC lines serve as vital genetic repositories for endangered suids. Our methodology provides a replicable framework for extending stem cell technology to other conservation-priority taxa, demonstrating how high-tech cellular tools can advance both fundamental research and biodiversity preservation against emerging pathogen threats.

## INTRODUCTION

Contemporary biodiversity faces unprecedented threats as species disappear at rates hundreds to thousands of times above natural background extinction levels, signaling the onset of Earth’s sixth mass extinction event^1,2^. This catastrophic biodiversity loss stems from multiple interconnected anthropogenic pressures including habitat destruction, overexploitation, invasive species, pollution, and climate change, all exacerbated by rapid human population growth and unsustainable consumption patterns^3^. Emerging infectious diseases now constitute an additional and increasingly significant driver of biodiversity decline, as pathogens spread among wildlife populations, cross species barriers, and destabilize ecosystem dynamics^4,5^. This convergence of ecological and disease pressures creates complex conservation challenges that demand innovative research approaches capable of addressing both immediate threats and long-term species preservation needs.

African swine fever (ASF) exemplifies this intersection between infectious disease and biodiversity conservation, representing one of the most devastating viral diseases affecting both domestic and wild suids globally^6,7^. Caused by ASF virus (ASFV), a complex double-stranded DNA virus of the family Asfarviridae, ASF exhibits remarkable environmental stability, sophisticated immune evasion mechanisms, and broad host range across phylogenetically diverse pig species^8,9^. The disease demonstrates striking species-specific pathogenicity patterns, with highly virulent strains causing acute hemorrhagic fever and near-100% mortality in domestic pigs and Eurasian wild boar (*Sus scrofa*) within days of infection, while African forest species such as Red river hogs (*Potamochoerus porcus*) and warthogs (*Phacochoerus africanus*) maintain asymptomatic infections despite cellular susceptibility^10,11^. This differential susceptibility landscape creates complex epidemiological dynamics that threaten vulnerable wild suid populations while simultaneously maintaining viral reservoirs in resistant species.

The global spread of ASF since 2007, from its endemic range in sub-Saharan Africa to Europe, Asia, and the Caribbean, has generated catastrophic impacts on both agricultural systems and wildlife conservation^12,13^. Economic losses exceed hundreds of billions of dollars globally, while ecological consequences include population crashes in susceptible wild boar populations and potential spillover risks to threatened endemic suids in affected regions^12,14^. The absence of commercially available vaccines or effective antiviral treatments, combined with ASFV’s environmental persistence and complex transmission cycles involving soft tick vectors, renders traditional control measures inadequate for protecting either domestic or wild populations^14,15^.

Current research approaches face substantial limitations that impede both mechanistic understanding and therapeutic development. Traditional *in vivo* studies using domestic pigs or wild boar are constrained by high costs, ethical considerations, biosafety requirements, and limited accessibility to threatened wild species^16^. Primary cell culture systems from wild suids present logistical challenges, particularly for endangered species where invasive sampling raises conservation concerns, while finite cell lifespan and phenotypic instability limit experimental scope^17^. These constraints are particularly problematic for comparative studies needed to understand species-specific susceptibility mechanisms and identify potential therapeutic targets based on natural resistance patterns.

Induced pluripotent stem cells (iPSCs) offer transformative opportunities to overcome these research limitations while simultaneously advancing conservation objectives^18,19^. iPSCs provide unlimited, renewable cell populations that can be differentiated into disease-relevant cell types, enabling standardized comparative studies across species without ongoing wildlife sampling requirements^20,21^. For endangered species, iPSCs represent a powerful *ex situ* conservation tool, preserving genetic diversity in a pluripotent state suitable for future applications including reproductive technologies and genetic rescue programs^22,23^.

This study presents the first successful integration-free reprogramming of somatic cells to iPSCs from four phylogenetically distinct porcine species representing critical nodes in the ASF susceptibility landscape: wild boar (*Sus scrofa*, Least Concern), Bornean bearded pig (*Sus barbatus*, Vulnerable), Babirusa (*Babyrousa babyrussa*, Vulnerable), and Red river hog (*Potamochoerus porcus,* Least Concern). These species span approximately 10 million years of evolutionary divergence and exhibit contrasting ASF susceptibility patterns, from high mortality in Asian *Sus* species to asymptomatic carriage in African *Potamochoerus*^11,24^. The successful derivation of integration-free iPSCs from these species, including two IUCN-listed threatened taxa, establishes an unprecedented cellular platform for comparative ASF research while creating valuable genetic resources for conservation applications.

Our approach employed Sendai virus-based reprogramming to ensure genomic integrity essential for both research validity and conservation applications, optimizing protocols for each species’ specific requirements. The resulting iPSC lines demonstrated robust pluripotency markers, tri-lineage differentiation capacity, and stable maintenance across extended passage, enabling their differentiation into CD14⁺CD11b⁺ monocytes - the primary ASFV target cells *in vivo*. This platform enables previously impossible comparative investigations of host-pathogen interactions across naturally resistant and susceptible species, while providing renewable genetic resources from threatened suids facing mounting conservation pressures.

## RESULTS

### Generation of Integration-Free Wild Boar ( *Sus scrofa* ) iPSCs

To establish pluripotent stem cells from Asian porcine species naturally susceptible to African swine fever (ASF), we derived primary fibroblasts from an ear biopsy of an adult wild boar female (Fig. 1a) and reprogrammed them using non-integrating Sendai virus vectors encoding the human transcription factors KLF4, OCT3/4, SOX2, and c-MYC (CytoTune™-iPS 2.0). Given that successful reprogramming of non-model species frequently requires species-tailored culture conditions^25^, we systematically screened multiple media formulations and extracellular matrix substrates to identify optimal parameters (Supplementary Fig. 1a).

**Figure 1.**
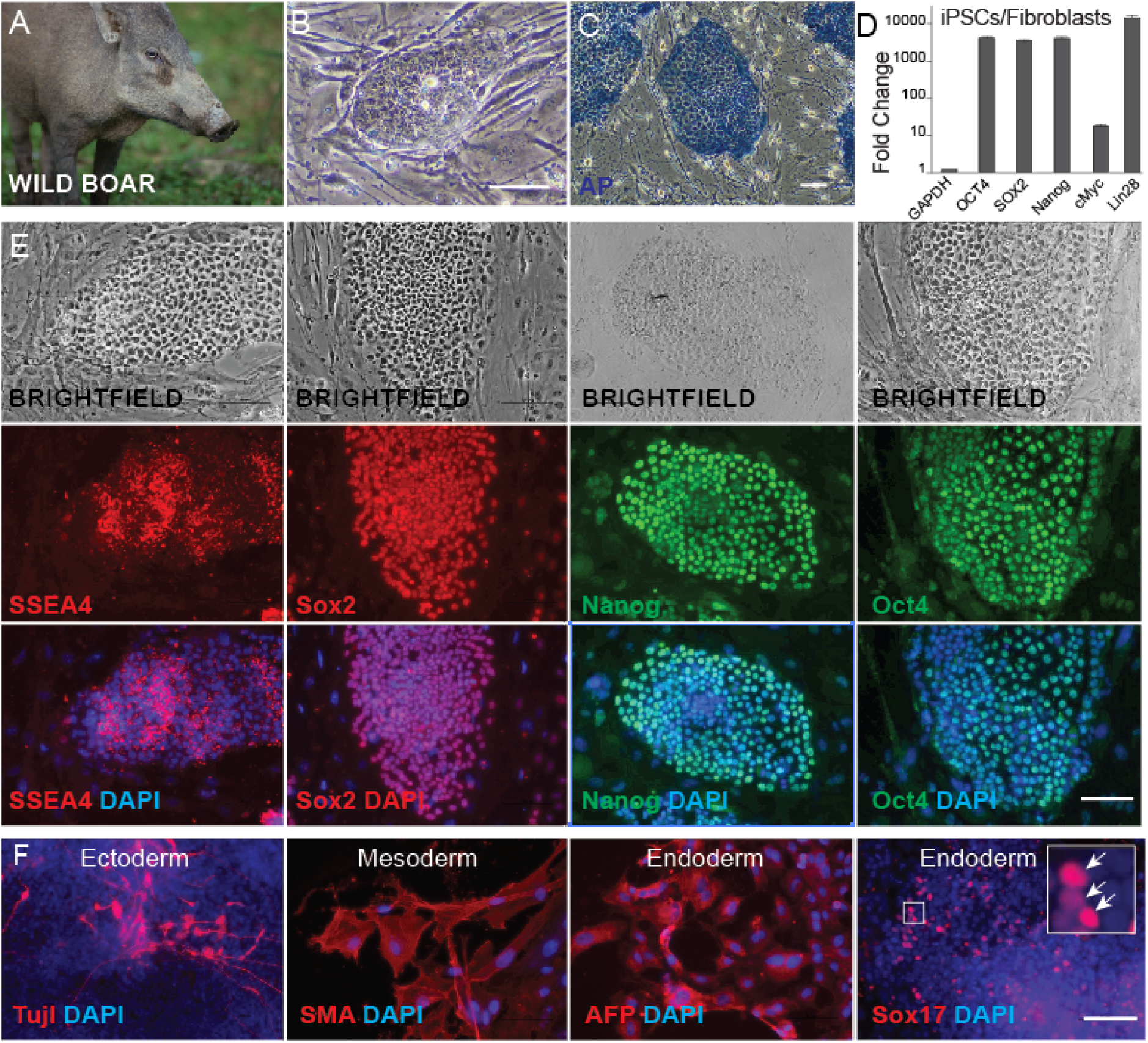
Generation and characterization of wild boar (*Sus scrofa* ) iPSCs. **(A)** Representative photograph of the Wild boar (*Sus scrofa*). **(B)** iPSC colony morphology at passage 3. **(C)** Alkaline phosphatase staining of iPSC^W^. **(D)** Quantitative RT-PCR analysis of endogenous pluripotency marker expression in iPSC^W^ relative to parental fibroblasts. **(E)** Immunofluorescence detection of SSEA4, SOX2, NANOG, and OCT4 in iPSC^W.^ **(F)** Tri-lineage differentiation capacity demonstrated by immunofluorescence staining for ectoderm (TUJ-1), mesoderm (α-SMA), and endoderm (AFP, SOX17) markers in day 7 embryoid bodies derived from iPSCW. Scale bars: 100 μm.

By day 5 post-transduction, morphological changes consistent with mesenchymal-to-epithelial transition were observed across all conditions tested. On day 8, cells were transferred to Matrigel- or LN521-coated six-well plates and cultured in porcine iPSC (piPS) medium supplemented with 1 µM A-83-01. Emerging colonies exhibited hallmarks of primed pluripotency, including flat, compact morphology with well-defined borders and a high nuclear-to-cytoplasmic ratio (Fig. 1b, c).

Reprogramming efficiency varied by medium composition. Pretreatment with Fibroblast medium (FM) supplemented with the TGF-β inhibitor A-83-01 yielded the highest frequency of alkaline phosphatase (AP)-positive colonies (0.26%), followed by FM with a small-molecule (SM) cocktail (0.21%), and N2B27 medium with SM (0.17%), as measured across independent replicates (Supplementary Fig. 1b). LN521 coating consistently supported higher reprogramming efficiency than Matrigel.

The resulting wild boar iPSC lines (iPSC^W^) were successfully propagated on irradiated mouse embryonic fibroblast (iMEF) feeders and maintained for a minimum of 15 passages without detectable loss of pluripotent identity. Transcriptional profiling and immunofluorescence (IF) staining confirmed robust endogenous expression of core pluripotency factors (Fig. 1d, e). To assess developmental potency, iPSC^W^ were cultured in bFGF- and hLIF-free hES medium, which triggered spontaneous embryoid body (EB) formation (Fig. 1f, Supplementary Fig. 1d). Following transfer to gelatine-coated dishes on day 7, the attached EBs differentiated into cells of all three germ layers, as confirmed by immunofluorescence staining for lineage-specific markers: TUJ-1 (ectoderm), α-SMA (mesoderm), and AFP and SOX17 (endoderm) (Fig. 1f). Collectively, these results demonstrate the successful generation of bona fide, integration-free wild boar iPSCs with broad developmental competence.

### Generation of Integration-Free Babirusa ( *Babyrousa babyrussa* ) iPSCs

To extend this reprogramming approach to threatened porcine species, we generated iPSCs from an 18-year-old female Babirusa (*Babyrousa babyrussa*) using the same Sendai virus-mediated strategy (Fig. 2a, b). Cells were cultured in FM for 6 days before transfer at day 7 onto LN521-, Matrigel-, or CF1 feeder-coated dishes (Supplementary Fig. 2a). From day 8, cultures were maintained in custom piPS medium with 1 µM A-83-01, either alone or in combination with the histone deacetylase inhibitor (HDACi) sodium butyrate.

**Figure 2.**
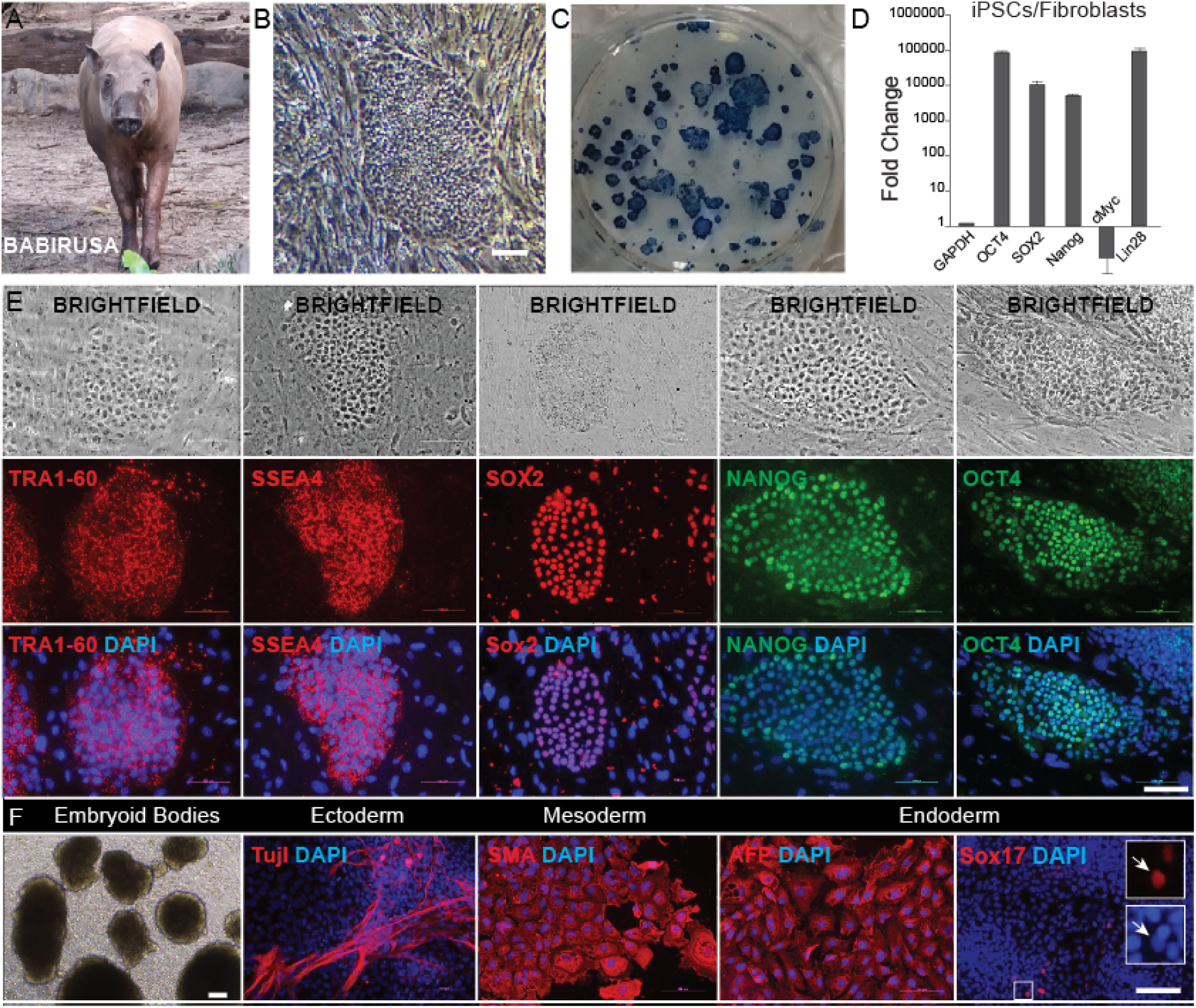
Generation and characterization of Babirusa (*Babyrousa babyrussa* ) iPSCs. **(A)** Representative photograph of the Babirusa (*Babyrousa babyrussa*). **(B)** iPSC^Ba^ colony morphology at passage 6. **(C)** Alkaline phosphatase staining of emerging iPSC^Ba^ colonies during reprogramming. **(D)** Quantitative RT-PCR analysis of endogenous pluripotency marker expression in iPSC^Ba^ relative to parental fibroblasts. **(E)** Immunofluorescence detection of TRA1-60, SSEA4, SOX2, NANOG, and OCT4 in iPSC^Ba^. **(F)** Tri-lineage differentiation capacity demonstrated by immunofluorescence staining for ectoderm (TUJ-1), mesoderm (α-SMA), and endoderm (AFP, SOX17) markers in day 14 embryoid bodies derived from iPSC^Ba^. Scale bars: 100 μm.

Primary colonies first appeared at day 20 post-transduction in piPS medium containing A-83-01, with characteristic pluripotent morphology established by day 30 (Fig. 2b, c). Colony emergence was strictly dependent on substrate, occurring exclusively on LN521-coated dishes at an efficiency of 0.12%, while reprogramming failed entirely on feeder layers and Matrigel (Supplementary Fig. 2b). The resulting Babirusa iPSC lines (iPSC^Ba^) maintained stable self-renewal through more than 15 passages on iMEF feeders in either piPS or mTeSR medium.

Pluripotent identity of iPSC^Ba^ was confirmed at the molecular level by elevated expression of endogenous pluripotency transcripts (Fig. 2d) and immunoreactivity for TRA1-60, SSEA4, SOX2, NANOG, and OCT4 (Fig. 2e). Functional pluripotency was demonstrated through spontaneous differentiation, with EBs forming across multiple passages (Fig. 2f) and, upon attachment, expressing lineage markers representative of all three germ layers: TUJ-1 (ectoderm), SMA (mesoderm), and AFP and SOX17 (endoderm). Collectively, these results confirm the successful derivation of bona fide pluripotent stem cells from Babirusa.

### Generation of integration-free Bornean bearded pig (Sus barbatus) iPSCs

Applying the same reprogramming strategy to an additional threatened porcine species, primary fibroblasts from a male Bornean bearded pig (Fig. 3a) were transduced using the CytoTune™-iPS 2.0 approach. Cells were cultured in FM with 10ng/ml bFGF for 8 days before transfer at day 7 onto Matrigel- or LN521-coated dishes, then maintained from day 8 in custom piPS medium supplemented with 1 µM A-83-01 (Supplementary Fig. 3a). Primary colonies with characteristic iPSC-like morphology emerged by day 12 post-transduction (Fig. 3b), at efficiencies of 0.013% on Matrigel and 0.02% on LN521. Established lines were maintained on iMEF feeders for more than 15 passages in mTeSR-plus medium; however, piPS medium was unable to support iPSC^Bp^ self-renewal.

**Figure 3.**
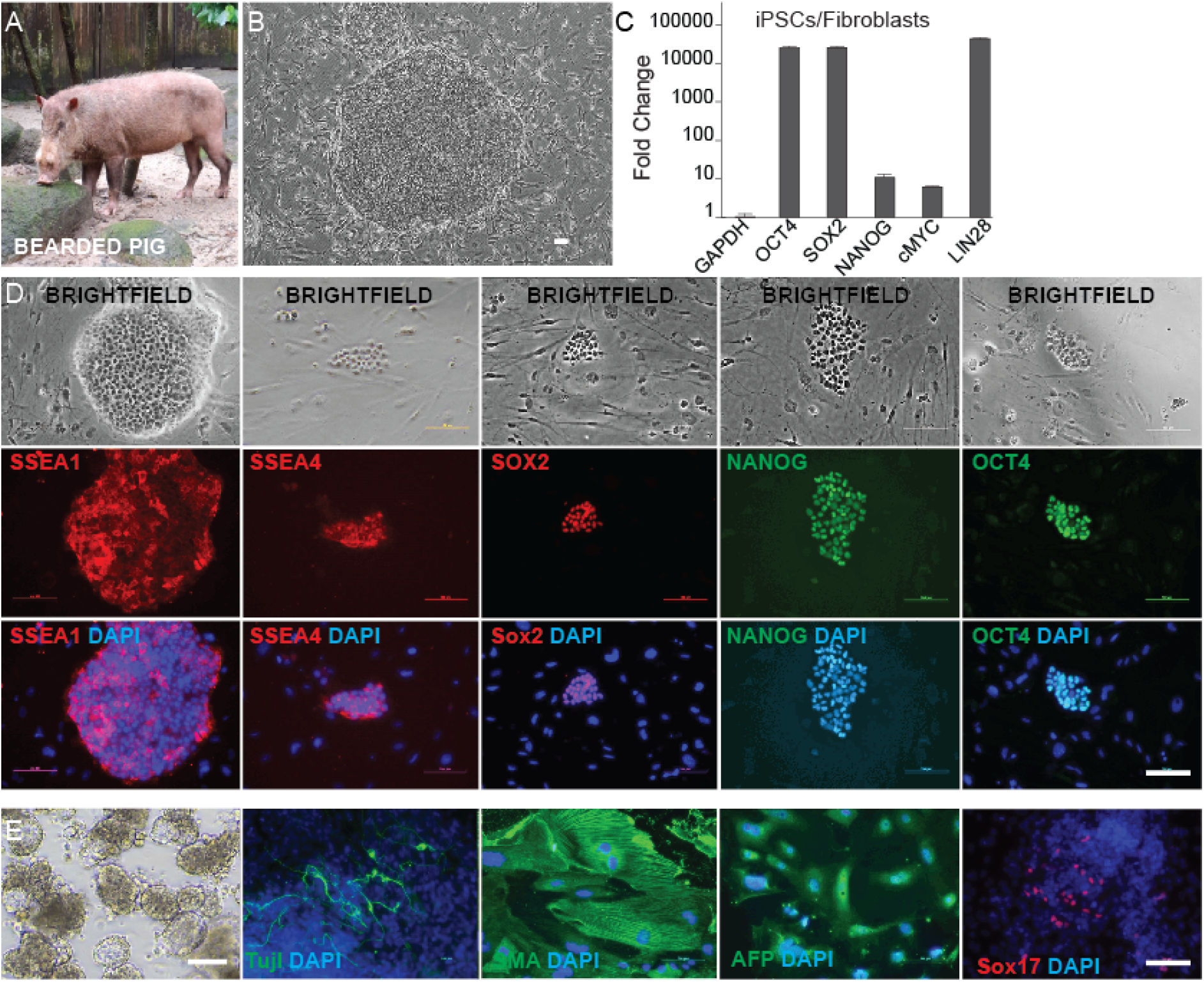
Derivation and characterization of integration-free Bornean bearded pig (*Sus barbatus* ) iPSCs. (A) Representative photograph of the Bornean bearded pig (*Sus barbatus*) donor. (B) Bright-field image of an iPSC^Bp^ colony at passage 6, illustrating characteristic pluripotent morphology with defined borders and high nuclear-to-cytoplasmic ratio. (C) Quantitative RT-PCR analysis of endogenous pluripotency gene expression (*OCT4*, *SOX2*, *NANOG*, *c-MYC*, *LIN28*) in iPSC^Bp^ relative to parental fibroblasts. (D) Immunofluorescence staining confirming nuclear and membrane localization of pluripotency-associated proteins SSEA1, SSEA4, SOX2, NANOG, and OCT4 in iPSC^Bp^. (E) Immunofluorescence staining of embryoid body outgrowths demonstrating tri-lineage differentiation capacity, with markers for ectoderm (TUJ-1), mesoderm (SMA), and endoderm (AFP and SOX17). Scale bars: 100 μm.

Molecular characterization confirmed successful reprogramming through significant upregulation of endogenous pluripotency transcripts, including OCT4, SOX2, NANOG, c-MYC, and LIN28, relative to the parental fibroblast population (Fig. 3c). IF further validated pluripotent identity through robust protein-level expression of SSEA1, SSEA4, SOX2, NANOG, and OCT4 (Fig. 3d). Notably, iPSC^Bp^ were negative for AP staining, pointing to species-specific differences in pluripotency marker expression compared to conventional human and mouse iPSC lines.

Functional pluripotency was assessed through spontaneous differentiation. Withdrawal of bFGF and hLIF prompted formation of three-dimensional EBs within 7 days across multiple passages (Fig. 3e). Transfer of day 7 EBs to gelatine-coated surfaces yielded attached outgrowths expressing lineage markers of all three germ layers: TUJ-1 (ectoderm), SMA (mesoderm), and SOX17 (endoderm) (Fig. 3e). Together, these findings confirm the successful derivation of functionally pluripotent stem cells from the Bornean bearded pig and highlight a notable species-specific divergence in AP expression.

### Generation of integration-free Red River Hog ( Potamochoerus porcus) iPSCs

To establish pluripotent stem cells from an African porcine species naturally resilient to ASF, PBMCs were isolated from a 12-year-old female Red river hog (Fig. 4a) and reprogrammed using the same CytoTune™-iPS 2.0 strategy. Cells were mixed with Sendai vector in the ultra-low attachment dishes in CD34+ medium, then topped the following day with custom piPS medium supplemented with 1 µM A-83-01, cells were seeded onto LN521 or feeder coated dishes in piPS media plus 1uM A-83-01 at day2 (Supplementary Fig. 4a). Colonies displaying morphological features consistent with mesenchymal-to-epithelial transition emerged by day 8 post-transduction in the LN521 coated wellat an efficiency of 0.003%, exhibiting positive AP staining (Fig. 4b). The resulting Red river hog iPSC lines (iPSC^Rrh^) maintained stable self-renewal through more than 15 passages on iMEF feeders in either piPS or mTeSR-plus medium.

**Figure 4.**
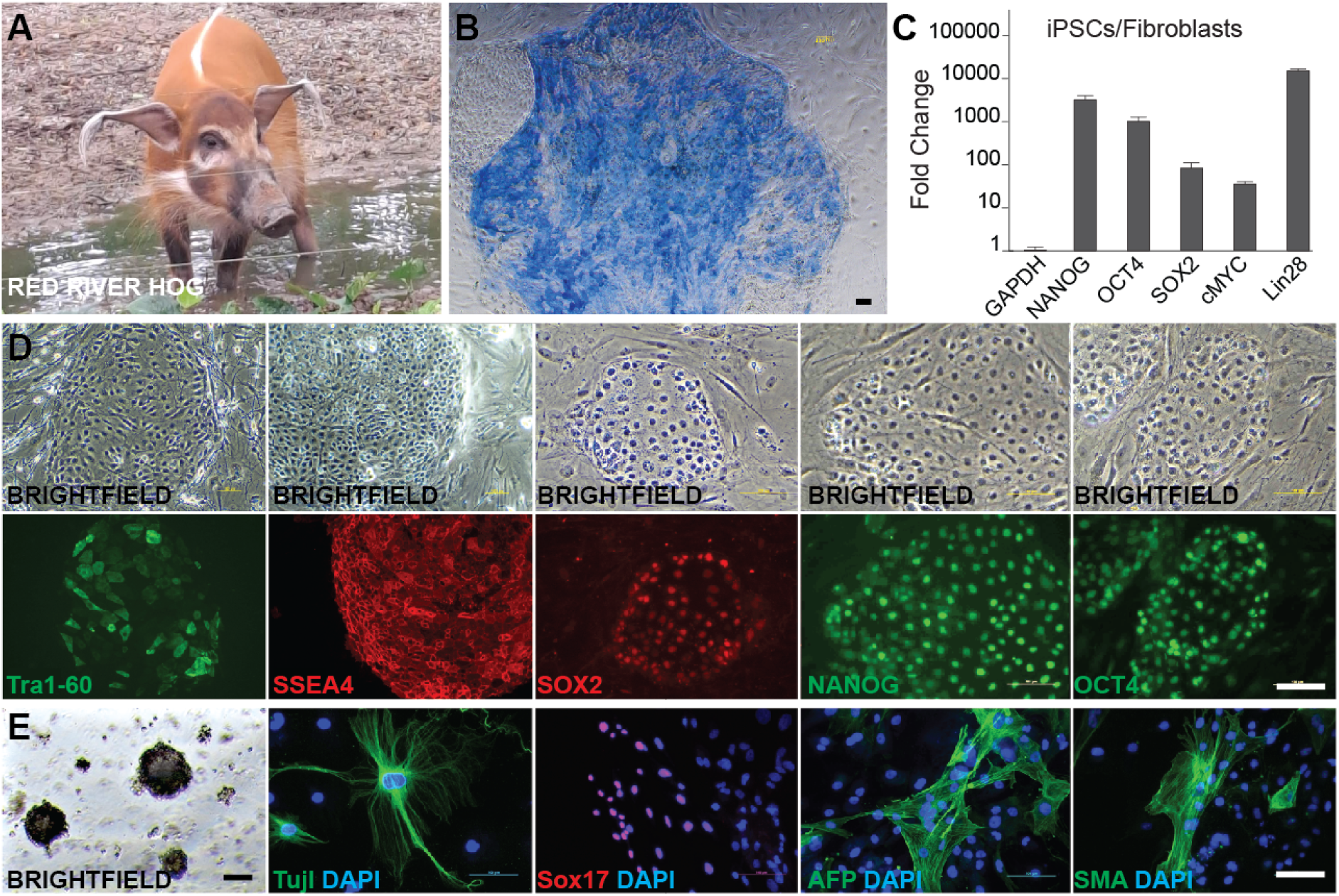
Derivation and characterization of integration-free Red River Hog (*Potamochoerus porcus* ) iPSCs. (A) Representative photograph of the Red River Hog (*Potamochoerus porcus*) donor. (B) Alkaline phosphatase staining of iPSC^Rrh^ colonies confirming pluripotent identity. (C) Quantitative RT-PCR analysis of endogenous pluripotency gene expression in iPSC^Rrh^ relative to parental fibroblasts. (D) Immunofluorescence staining for TRA1-60, SSEA4, SOX2, NANOG, and OCT4, confirming pluripotency marker expression in iPSC^Rrh^. (E) Immunofluorescence staining of embryoid body outgrowths demonstrating tri-lineage differentiation capacity, with markers for ectoderm (TUJ-1), mesoderm (SMA), and endoderm (AFP and SOX17). Scale bars: 100 μm.

Pluripotent identity was confirmed at the molecular level by elevated expression of endogenous pluripotency transcripts (Fig. 4c) and immunoreactivity for TRA1-60, SSEA4, SOX2, NANOG, and OCT4 (Fig. 4d). Functional pluripotency was demonstrated through spontaneous differentiation, with EBs forming across multiple passages (Fig. 4e) and, upon attachment, expressing lineage markers representative of all three germ layers: TUJ-1 (ectoderm), SMA (mesoderm), and AFP and SOX17 (endoderm). Collectively, these results confirm the successful derivation of bona fide pluripotent stem cells from the Red river hog.

### Generation of ASF-relevant monocytes from iPSCs of four porcine species

To develop a comparative in vitro model system for ASF research, we established monocyte differentiation protocols using iPSCs derived from four phylogenetically distinct porcine species with varying ASF susceptibility profiles. Our species panel included three ASF-susceptible species (Babirusa, Bearded pig, and Wild boar) and one natural asymptomatic carrier species (Red river hog).

iPSCs from each species were subjected to a standardized three-stage differentiation protocol conducted in 2D adherent cultures ((Supplementary Fig. 5a). This stepwise approach progressed through sequential phases: mesodermal commitment, hematopoietic specification, and terminal monocyte differentiation. The protocol successfully generated CD14⁺CD11b⁺ monocytes from iPSC^W^ (Wild boar), iPSC^Ba^ (Babirusa), and iPSC^Rrh^ (Red river hog) cell lines.

Monocyte populations emerged in culture supernatants as early as day 14 post-differentiation initiation, enabling continuous harvesting throughout the remainder of the culture period to optimize cell yield. IF analysis confirmed robust monocyte differentiation across successfully differentiated cell lines, with the proportion of CD14⁺CD11b⁺ double-positive cells exceeding 60% in populations derived from Wild boar, Babirusa, and Red river hog iPSCs (Fig. 5). However, significant inter-species variations in differentiation success were observed. iPSC^Bp^ (Bearded pig) cells demonstrated poor yield during the differentiation process, with most cells failing to complete terminal monocyte differentiation (Supplementary Fig. 5b). Also, iPSC^Rrh^ exhibited reduced viability compared to iPSC^W^ and iPSC^Ba^, resulting in lower overall cell yields, though surviving cells showed comparable differentiation efficiency (>60%). Thus, this yield limitation can be readily addressed by initiating differentiation with larger numbers of starting iPSCs. Notably, the CD14⁺CD11b⁺ cells displayed heterogeneous expression patterns, with some cells exhibiting strong expression while others showed weaker expression of these markers (Fig. 5), potentially reflecting different maturation states within the monocyte population or species-specific variations in marker expression intensity.

**Figure 5.**
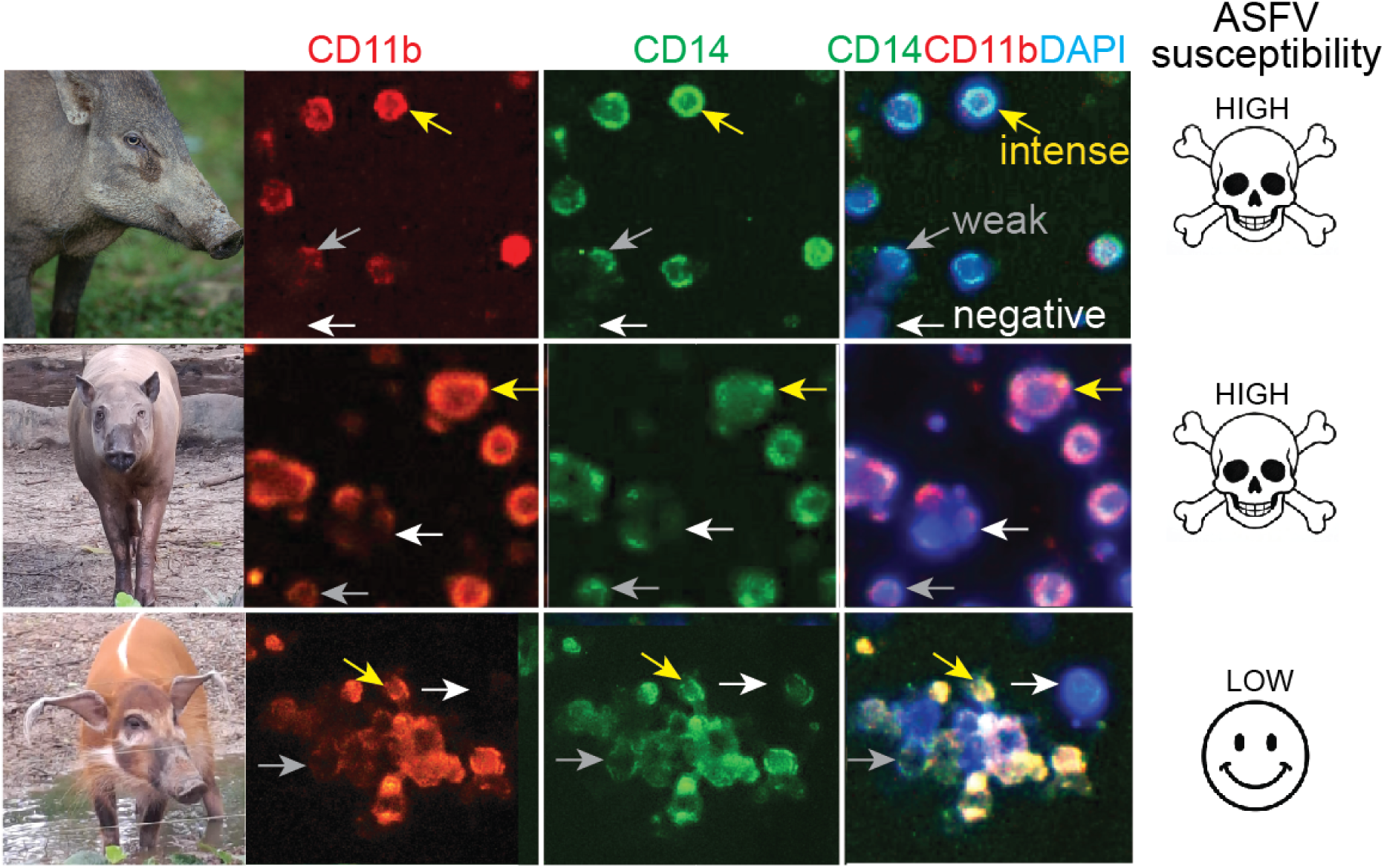
Monocyte differentiation from wild boar, Babirusa, and Red River Hog iPSCs. Immunofluorescence staining for CD14 and CD11b expression in monocytes differentiated from iPSC^W^ (wild boar), iPSC^Ba^ (Babirusa), and iPSC^Rrh^ (Red River Hog), complementing the data shown in Figure 5. Yellow arrows indicate monocyte-like cells with high CD14 and CD11b expression; grey arrows indicate cells with lower marker expression. White arrows denote CD14⁻CD11b⁻ cells, which constitute fewer than 40% of the total population. Scale bars: 100 μm.

These findings demonstrate the successful establishment of a multi-species iPSC-derived monocyte differentiation platform that recapitulates the natural diversity of ASF susceptibility observed across porcine species. The observed inter-species differences in differentiation efficiency, cell survival, and marker expression patterns likely reflect intrinsic biological variations between species, providing a valuable framework for comparative ASF pathogenesis studies. This platform enables investigation of species-specific responses to ASFV infection using the primary target cell type in a controlled in vitro environment.

## DISCUSSION

This study represents the first comprehensive study establishing integration-free induced pluripotent stem cells (iPSCs) from four phylogenetically distinct porcine species: wild boar (Sus scrofa), Babirusa (Babyrousa babyrussa), Bornean bearded pig (Sus barbatus), and Red river hog (Potamochoerus porcus). These species occupy critical positions along the African swine fever virus (ASFV) susceptibility spectrum, with Red river hogs demonstrating natural asymptomatic carriage while Asian suids exhibit severe mortality upon infection ^11,16^. The successful derivation of these iPSC lines establishes an unprecedented cellular platform for comparative ASF research and conservation applications in endangered suid species.

Reprogramming efficiency demonstrated substantial interspecies variation, ranging from 0.003% to 0.26%, consistent with documented differences in pluripotency induction even across related mammalian species^26^. Wild boar fibroblasts achieved optimal efficiency (0.26%), comparable to established domestic pig iPSC protocols^20,27^, while Red river hog peripheral blood mononuclear cells (PBMCs) exhibited minimal efficiency (0.003%). This disparity likely reflects both cell-type-specific reprogramming barriers and species-intrinsic factors affecting pluripotency induction^28,29^. Intermediate efficiencies observed in Babirusa (0.12%) and Bornean bearded pig (0.02%) suggest a continuum of reprogramming competence across the suid lineage. These efficiency variations potentially may arise from differential endogenous expression of Yamanaka factors (OCT4, SOX2, KLF4, c-MYC), species-specific epigenetic landscapes, or varying requirements for mesenchymal-to-epithelial transition signaling pathways^30,31^.

Culture condition optimization revealed species-specific requirements across all derivations. TGF-β pathway inhibition via A-83-01 proved universally essential, confirming its established role in maintaining naive/primed pluripotency states and preventing spontaneous differentiation^32^. However, substrate dependencies exhibited remarkable species specificity. Babirusa reprogramming demonstrated absolute requirement for LN521-coated surfaces, failing completely on Matrigel or feeder layers. This stringent substrate selectivity suggests divergent integrin-mediated adhesion signaling or extracellular matrix interaction requirements during pluripotency establishment^33,34^. Similar substrate specificities have been documented in non-model species iPSC derivations, emphasizing the critical importance of systematic protocol optimization beyond established model organisms^19,35^.

A particularly intriguing observation was the absence of alkaline phosphatase (AP) activity in Bornean bearded pig iPSCs despite robust expression of core pluripotency transcription factors (OCT4, SOX2, NANOG) and demonstrated tri-lineage differentiation competence. This phenotype parallels findings in porcine epiblast stem cells, where AP negativity does not preclude authentic pluripotency^36,37^. The molecular mechanisms underlying this divergence remain unclear but may involve species-specific *ALPL* gene regulation, post-translational modifications, or alternative alkaline phosphatase isoform expression patterns^38,39^. This finding underscores the limitations of single-marker pluripotency validation across phylogenetically diverse species and emphasizes the necessity for comprehensive molecular and functional characterization approaches.

Medium compatibility analysis revealed further evidence of species-specific culture requirements that extend beyond reprogramming protocols to maintenance conditions. While all four iPSC lines demonstrated stable self-renewal and pluripotency marker expression when cultured on inactivated mouse embryonic fibroblast (iMEF) feeders in mTeSR-plus medium, transition to defined, feeder-free conditions exposed pronounced differential responses. The optimized piPS successfully supported long-term culture of Red river hog, Babirusa, and wild boar iPSCs while completely failing to maintain Bornean bearded pig iPSCs, which underwent rapid differentiation. This selective medium incompatibility likely reflects interspecies variation in growth factor receptor expression profiles, downstream signalling pathway dependencies, or fundamental metabolic requirements that have diverged during suid evolution^34^. These findings have significant practical implications for establishing robust culture systems when extending iPSC technology to non-model species, highlighting the necessity for systematic medium optimization rather than assuming universal applicability of protocols developed for domestic or laboratory animals. Such medium selectivity considerations become particularly critical for conservation applications where maintaining genetic integrity across extended culture periods is essential for downstream applications.

### iPSC-Derived Monocytes as an ASF Research Platform

The successful differentiation of all four porcine iPSC lines into CD14⁺CD11b⁺ monocytes establishes a renewable, species-comparative research platform addressing critical limitations in ASF pathogenesis studies. Peak differentiation efficiency exceeded 60% across most species, demonstrating robust myeloid lineage specification from primed pluripotent cells. This efficiency compares favourably with reported human and murine iPSC-derived myeloid differentiation protocols, suggesting conservation of core hematopoietic developmental programs across suids despite ∼5-10 million years of evolutionary divergence.

Despite consistent differentiation efficiency, substantial interspecies variation in absolute cell yield was observed, with wild boar and Babirusa iPSCs producing significantly higher cell numbers than Red river hog and Bornean bearded pig lines. This differential proliferative capacity may reflect intrinsic differences in hematopoietic progenitor expansion potential, species-specific metabolic requirements during differentiation, or suboptimal culture conditions requiring further optimization.

This iPSC-monocyte system addresses fundamental limitations in ASF research infrastructure. Primary monocyte/macrophage isolation from wild suids presents significant logistical challenges, particularly for threatened species like Babirusa and Bornean bearded pig (both classified as Vulnerable by IUCN), where even sample collection from non-threatened species raises ethical concerns. Additionally, primary cultures suffer from limited lifespan, phenotypic heterogeneity, and donor-dependent variation that confound reproducible experimental design. iPSC-derived monocytes circumvent these constraints by providing unlimited, genetically defined cell populations amenable to standardized protocols, high-throughput screening, and genetic manipulation approaches.

This platform enables previously unattainable research directions informed by recent breakthroughs in ASFV infection mechanisms. Comparative infection studies using genetically matched monocytes from susceptible versus resistant species can systematically identify host determinants of viral tropism, replication efficiency, and immune evasion mechanisms. The recent discovery that ASFV entry requires synergistic cooperation between CD163, CD169/Siglec1, and the newly identified co-receptor MYH9 (non-muscle myosin heavy chain 9) provides specific targets for comparative analysis across species^40,41^. CRISPR-mediated genome editing in iPSCs followed by monocyte differentiation permits functional validation of candidate resistance alleles identified through comparative genomics. Species-specific polymorphisms in RELA (NF-κB signaling) and MX1 (interferon-induced antiviral responses) were also identified as potential ASF resistance determinants in African suids^42^. These findings gain new significance given the extensive ASFV targeting of the cGAS-STING pathway, where over 20 viral proteins systematically suppress type I interferon production through multiple mechanisms including STING degradation (MGF505-6R), IRF9 sequestration (MGF505-7R), TBK1 degradation (MGF505-3R), and novel ferroptosis-mediated immune suppression (A151R)^41,43–45^. iPSC-derived monocytes provide a tractable experimental system to test whether introducing resistance-associated alleles into susceptible species’ iPSCs confers protective phenotypes, while also enabling investigation of species-specific susceptibility to ASFV’s sophisticated immune evasion strategies. Furthermore, this platform allows systematic evaluation of the recently discovered AXL receptor-mediated apoptotic mimicry entry mechanism and ferroptosis-induced metabolic reprogramming that ASFV exploits for infection and persistence^41,45^.

Furthermore, this system facilitates mechanistic dissection of ASFV entry, replication cycles, and innate immune signalling in controlled, renewable cell populations. The ability to generate monocytes from diverse genetic backgrounds enables identification of conserved versus species-specific host-pathogen interactions, critical for developing broad-spectrum antiviral therapeutics or species-targeted intervention strategies^46,47^. For threatened species conservation, iPSC-derived monocytes enable ASF research without requiring ongoing wildlife sampling, aligning with conservation objectives while advancing pathogen understanding.

### Limitations and Future Directions

Several important limitations require consideration. First, while iPSC-derived monocytes express appropriate phenotypic markers (CD14, CD11b), their functional equivalence to primary tissue-resident macrophages - the principal ASFV targets in vivo - requires comprehensive validation. Transcriptomic and proteomic profiling comparing iPSC-derived cells to ex vivo monocyte-derived macrophages would clarify whether developmental origin influences infection susceptibility, innate immune activation, or viral replication kinetics. Studies in human iPSC systems indicate that iPSC-derived monocytic cells are functionally comparable to primary cells, successfully mimicking cytokine profiles and core surface markers, which makes them robust tools for disease modelling^48,49^. However, they are not identical - they exhibit lower HLA-DR expression, higher IL-10 levels, and variability between different iPSC lines which may reflect incomplete maturation states.

Second, observed interspecies variation in monocyte yield may reflect suboptimal culture conditions rather than intrinsic biological differences. Systematic optimization of cytokine combinations, small molecule supplements, and temporal differentiation parameters for each species could improve yields and potentially reduce performance disparities. Species-specific serum formulations or three-dimensional culture systems may better support hematopoietic differentiation in lower-performing lines^50,51^.

Finally, current monocyte differentiation protocols yield heterogeneous populations. While CD14⁺CD11b⁺ cells comprise the predominant fraction (>60%), single-cell RNA sequencing would resolve whether subpopulations recapitulate in vivo monocyte diversity (classical, intermediate, non-classical subsets) and whether species exhibit differential subset compositions^52^. Such cellular heterogeneity may prove critical, as ASFV tropism can demonstrate subset specificity^47^.

### Broader Implications

Beyond ASF research applications, the establishment of these iPSC lines creates the foundation for a dedicated conservation biobank that can serve global research communities working at the intersection of conservation biology, comparative genomics, and disease ecology. Babirusa and Bornean bearded pig populations face severe threats from habitat destruction and hunting pressure, with both species classified as Vulnerable by IUCN assessments. iPSCs from these species constitute genetic repositories for potential future applications including genome rescue strategies (introducing genetic diversity from archived cell lines into declining populations), disease modeling for additional emerging pathogen threats, and assisted reproductive technologies^19,53^.

From evolutionary perspectives, this suid iPSC panel offers unique opportunities to investigate pluripotency network evolution across mammalian radiations. Comparative transcriptomic and epigenomic analyses could reveal lineage-specific adaptations in core regulatory circuits, identify conserved versus divergent enhancer elements governing pluripotency gene expression, and elucidate how developmental plasticity mechanisms have evolved across phylogenetic scales^54,55^. The observed phenotypic variations (AP negativity, substrate selectivity, medium dependencies) likely reflect deeper molecular divergences that, once characterized, could inform optimization strategies for additional non-model species.

## Conclusion s

This study successfully establishes integration-free iPSCs from four evolutionarily distinct porcine species representing critical nodes in the ASFV susceptibility landscape and demonstrates their robust differentiation into monocytes, the primary viral target cells. The resulting experimental platform enables comparative host-pathogen investigations previously constrained by limited access to wild suid primary cells, particularly from conservation-priority species. Documented interspecies variation in reprogramming efficiency, culture requirements, and monocyte production capacity highlights both challenges and opportunities inherent in extending stem cell technologies beyond traditional model organisms.

Future research priorities include validating functional equivalence between iPSC-derived and primary macrophages, optimizing species-specific culture protocols, and conducting systematic comparative ASFV infection studies. These efforts will ultimately determine the platform’s utility for understanding and mitigating ASF, a devastating disease affecting global swine populations, biodiversity conservation, and food security worldwide.

## MATERIALS AND METHODS

### Ethics statement

This project has been approved by the MWG Research Panel under the project code MWG230104 and MWG250403. No animals were harmed in the preparation of this manuscript

### Resource availability

This study generated new unique resources. Further information and requests for resources and reagents should be directed to the corresponding author from Mandai Nature.

### Access to skin samples and animal used for this study

The ESCAR laboratory is part of the Mandai Wildlife Group, which is the steward of Mandai Wildlife Reserve, home to Singapore Zoo, Night Safari, River Wonders, and Bird Paradise. Animals either died of natural causes or were humanely euthanised due to medical reasons. Skin samples were obtained from only one donor animal per species.

### Animal details

#### Babirusa (Babyrousa babyrussa)

Female (ZIMS ID: G9234), born in captivity at Singapore Zoological Gardens on 23 October 2002. The animal died on 6 July 2021 at 18 years of age; cause of death was undetermined. Skin was collected at post-mortem for fibroblast derivation.

#### Red river hog (Potamochoerus porcus)

Female (ZIMS ID: G11823), born in captivity at Singapore Zoological Gardens on 26 April 2010. Peripheral blood was collected for reprogramming prior to the animal’s transfer to Avilon Zoo, Philippines. The animal was 11 years of age at the time of sampling.

#### Bornean bearded pig (*Sus barbatus*)

Male (ZIMS ID: G19357; microchip: 00-07E0-53E5), born in captivity on 1 January 2022. Skin was collected for fibroblast derivation in 2022. At the time of submission, this individual remains alive and is housed at Singapore Zoological Gardens.

#### Wild boar (*Sus scrofa*)

Male of unknown age, received from the wild. Skin was collected in 2019 for fibroblast derivation.

### Derivation and culture of porcine primary fibroblasts

At post-mortem, the animals’ skin was shaved before aseptic surgical preparation of the area of interest with chlorhexidine and 70% ethanol. A sterile scalpel blade was used to harvest a 3 cm x 3 cm sample of full-thickness skin. In sterile conditions, any remaining fur, fat, and epidermis were removed from the skin sample, and the sample was cut into smaller pieces. After washing in PBS, the pieces were incubated in DMEM, 1X Antibiotic–Antimycotic, and 0.5 mg/ml Fungin™, at room temperature for 30 min. After further washing in PBS, the pieces were placed in a 0.1% gelatin-coated sterile tissue culture dishe with media containing FM-p0, and incubated at 37 degrees Celsius and 5% CO2. When confluent, fibroblasts were passaged with 0.25% trypsin and maintained on 0.1% gelatine-coated sterile tissue culture dishes with FM. For efficient reprogramming, fibroblasts were transduced or transfected at passage 5 or earlier

### Medium compositions

N2B27 media: 50% Neurobasal, 50% DMEM/F12 (1:1), 0.5X N-2 Supplement and 0.5 X B27 Supplement, 100 uM β-mercaptoethanol, 2mM L-Glutamine, 1X Penicillin–Streptomycin

SM cocktail: 10uM Y-27632, 0.5uM PD0325901, 3uM CHIR99021, 0.5uM A-83-01, 10uM Forskolin

Fibroblast media (FM): DMEM, 10% FBS, 1X MEM NEAA, 100 µM β-mercaptoethanol, 1X Penicillin–Streptomycin, and 2 mM L-glutamine

piPS media: DMEM/F12; 15% ES FBS, 1X MEM nonessential amino acids (NEAA), 100 uM β-mercaptoethanol; 1X penicillin/ streptomycin; 2 mM L-glutamine; 10 ng/ml bFGF

hES media; piPS media+10ng/ml hLIF

FM-p0 media: DMEM, 20% FBS, 1X MEM non-essential amino acids (NEAA), 1X Penicillin–Streptomycin, and 2 mM L-glutamine

### The isolation and culture of red river hog blood cells

Whole blood was diluted1:1 with PBS before it was layered gently over Ficoll-Paque (2 times the volume of diluted blood) in 50ml falcon tube. The tube was centrifuged at 450x g for 30 mins at room temperature with brake off. The white layer of cells (PBMC) at the interphase were collected carefully and transferred into a new falcon tube. The cells were washed with PBS twice (5-10 volumes of cells) and centrifuged at 350xg for 10 mins with brake on. If the pellet (PBMC) was red, the pellet was resuspended with 10ml red blood lysis buffer (1:10 dilution) and incubated at RT for 10mins. After centrifugation at 350xg for 10 mins, the clean PBMC were resuspended into culture media for expansion or freezing media for cryopreservation

### Derivation and culture of Babirusa iPSCs

300K cells were seeded into 3cm dishes, cultured in FM; 24 hours later (day0), the cells were infected with CytoTune™ Sendai vector containing human Klf4, Oct3/4, Sox2 and C-Myc in 1ml FM. Next day (day1), the cells were cultured in the FM containing 10ng/ml bFGF. The media were changed daily. On day7, the cells were passaged and seeded into 6 well plates coated with irradiated CF1 mouse embryonic fibroblast (iMEF) feeder layers, Matrigel or LN521 as 1:4 ratio and cultured with FM with bFGF. On day 8, the media were changed to piPS media containing DMEM/F12; 15% ES FBS, 1X MEM nonessential amino acids (NEAA), 100 uM β-mercaptoethanol; 1X penicillin/ streptomycin; 2 mM L-glutamine; 10 ng/ml bFGF, plus extra 1uM A083. Colonies with typical iPSCs-like morphology appeared from day 20. Cells grown on LN512 without the Histone Deacetylase (HDAC) inhibitor sodium butyrate showed significantly better reprogramming efficiency compared with the other conditions. These colonies were allowed to expand in size. After 5-14 days, colonies were mechanically picked and expanded in 96 well plate seeded with feeders in piPS media. The cells maintained their characteristic morphology, were stable, and could be passaged more than 15 passages (further passaging have not been tested).

### Derivation and culture of Wild Boar iPSCs

300k fibroblast cells were seeded into 3 wells of 12 well plate, cultured in FM; 24 hours later (day0), the cells were infected with CytoTune™ Sendai vector containing human Klf4, Oct3/4, Sox2 and C-Myc in 1.2ml FM. Next day (day1), the cells were cultured in the FM containing 10ng/ml bFGF. The media were changed to N2B27 media (see materials and methods) +10ng/ml bFGF+SM cocktail; FM + 10ng/ml bFGF+SM cocktail; or FM + 1uM A83-01. On day 8, the cells were passaged into Matrigel coated or LN521 coated 6 well plate, cultured in piPS media plus 1uM A083. The colonies were picked and expanded in 96 well plate coated with feeder cells at day 23. Cell morphology changed 5 days post transduction. Reprogramming was most efficient in N2B27 media with SM cocktail, least in FM plus A-83-01 when the reprogramming started. However, the final reprogramming efficiency showed opposite result

### Reprogramming of Red river hog blood cells

PBMC were isolated from whole blood with Ficoll and cultured in CD34+ media (SFEMII plus commercial cocktail) for 8 days. At day 0, 100K cells in 400ul CD34+ media were infected with CytoTune™ Sendai vector containing human Klf4, Oct3/4, Sox2 and C-Myc in ultralow attachment 24 well plate. Next day, 2.6 ml piPS media + 1uM A-83-01 were added into the well (day1). 24 hrs later, the cells were split into two wells coated with feeder cells or LN521 cultured in piPS media + 1uM A-83-01 (day2). The media were changed every two days. At day8, there are reprogrammed cells attached to the bottom of the LN521 coated well. 1 month later, 3 colonies were picked and expanded in 48 well plate coated with feeder cells in piPS media, only 2 colonies were successfully expanded.

### Reprogramming of bearded pig fibroblast cells

300K cells were seeded into 3cm dishes, cultured in FM; 24 hours later (day0), the cells were infected with CytoTune™ Sendai vector containing human Klf4, Oct3/4, Sox2 and C-Myc in 1ml FM plus 20ng/ml bFGF. Next day (day1), the cells were cultured in 2ml fresh FM containing 20ng/ml bFGF and 1uM A-83-01. At day6, the cells were passaged and seeded into 6 well plates coated with Matrigel or LN521 in FM containing 15ng/ml bFGF and 1uM A-83-01. At day7, the cells were cultured in piPS media with 1uM A-83-01. The media were changed every two days. The colonies were detected in the reprogramming well around day12. On day 20, colonies were picked and expanded into 96 well plate coated with LN521 in mTeSR-plus media. The iPSC needs to be cultured into the wells coated with feeder cells in mTeSR-plus to maintain pluripotent state after P4.

### Embryoid body formation and differentiation into three germ layers

iPSCs were passaged with ReLeSR™ (STEMCELL Technologies, #100–0484) and split 1:3 into ultralow attachment 24-well plates. Cells were cultured in hES media without bFGF and hLIF for 7 days. EBs formed were either maintained in suspension for another 7 days before being harvested for RNA extraction or transferred to a 0.1% gelatin-coated plate for another 7 days before being fixed for immunocytochemical analysis.

## ACKNOWLEDGEMENTS

This paper is a tribute to the late Dr. Chai CHOU, whose pioneering spirit and dedication as our former lab head continue to guide our research. His presence is greatly missed. We also thank David Tan, Senior photographer an Mandai Wildlife Group for providing the Wild boar’s picture.

## AUTHOR CONTRIBUTIONS STATEMENT

Conception and design: Q.B.(equal), O.P.(equal); Methodology, Analysis and interpretation of data, Investigation: Q.B (lead), C.Y.L. (supporting), N.L.T. (supporting), H.L.Y. (supporting), K.P. (supporting), C.H. (supporting), S.M.C. (supporting); Writing and revising the article, O.P. (lead), Q.B. (supporting), C.Y.L. (supporting), N.L.T. (supporting); Supervision: O.P. (lead), S.C.N. (supporting), Y.H.L. (supporting), S.X. (supporting); Funding acquisition: S.C.N. (lead), N.L.T. (supporting).

## SUPPLEMANTRY INFORMATION

**Supplementary Figure 1.**
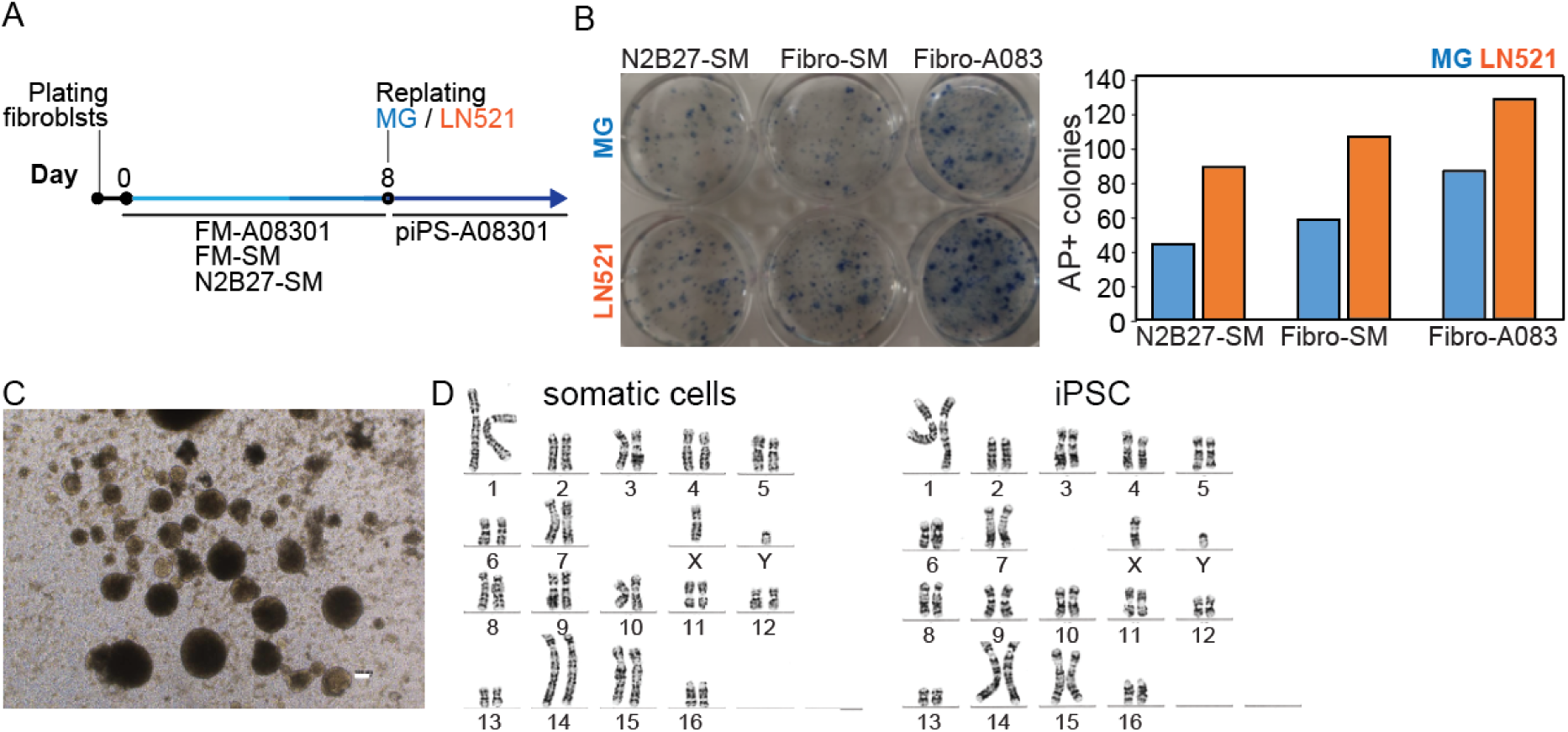
Reprogramming strategy and characterization of wild boar (*Sus scrofa* ) iPSCs. (A) Schematic overview of the Sendai virus-mediated reprogramming protocol for wild boar fibroblasts. (B) Alkaline phosphatase staining of iPSC^W^ colonies in reprogramming wells at day 25 (left) and quantification of colony yield under different culture conditions (right). (C) Representative images of embryoid body formation from iPSC^W^. (D) Karyotype analysis of parental fibroblasts and iPSC^W^ (38, XY).

**Supplementary Figure 2.**
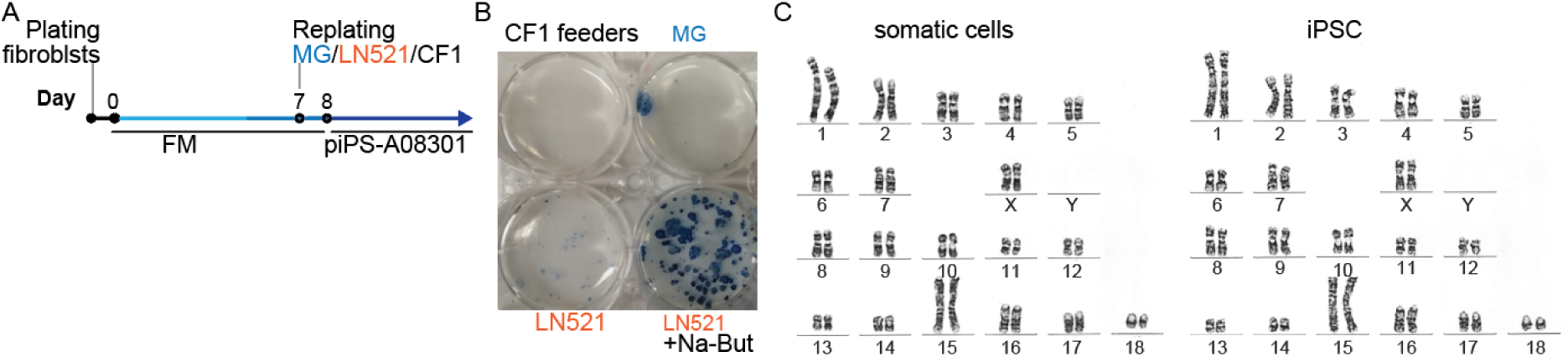
Reprogramming strategy and characterization of Babirusa (*Babyrousa babyrussa*) iPSCs. (A) Schematic overview of the Sendai virus-mediated reprogramming protocol for Babirusa fibroblasts. (B) Alkaline phosphatase staining of iPSC^Ba^ colonies in the original reprogramming plate. (C) Karyotype analysis of parental fibroblasts and iPSC^Ba^, confirming a normal female karyotype (38, XX).

**Supplementary Figure 3.**
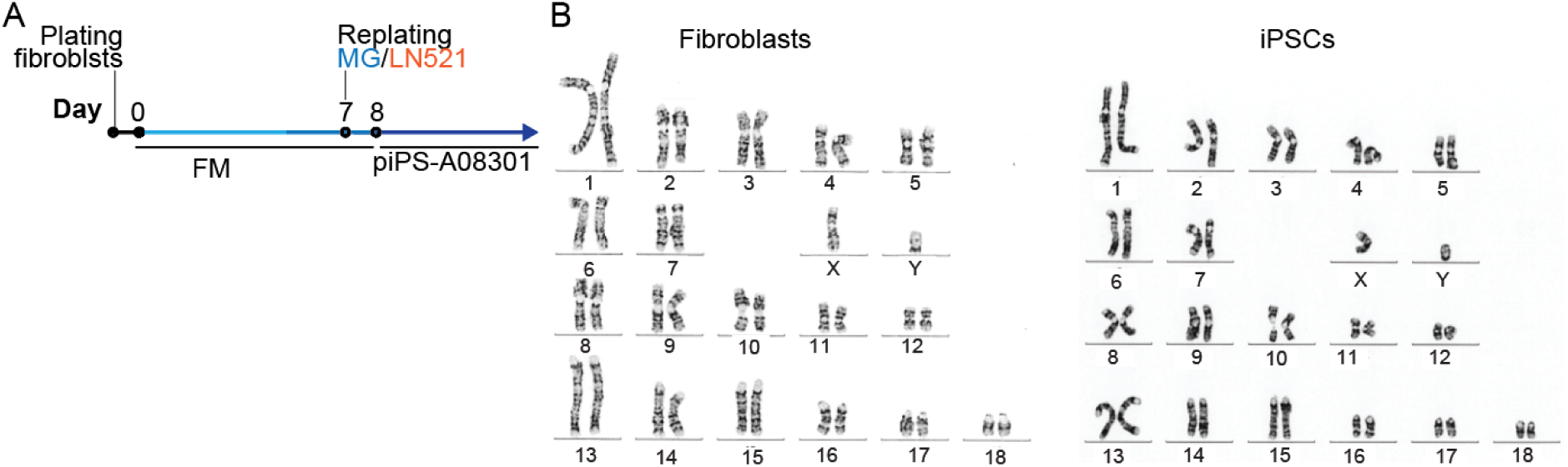
Reprogramming strategy and characterization of Bornean bearded pig (*Sus barbatus*) iPSCs. (A) Schematic timeline of the Sendai virus-mediated reprogramming protocol for Bornean bearded pig fibroblasts. (B) RT-PCR analysis confirming complete clearance of Sendai virus vectors from established iPSC^Bp^ lines. (C) Karyotype analysis of parental fibroblasts and iPSC^Bp^, confirming a normal male karyotype (38, XY).

**Supplementary Figure 4.**
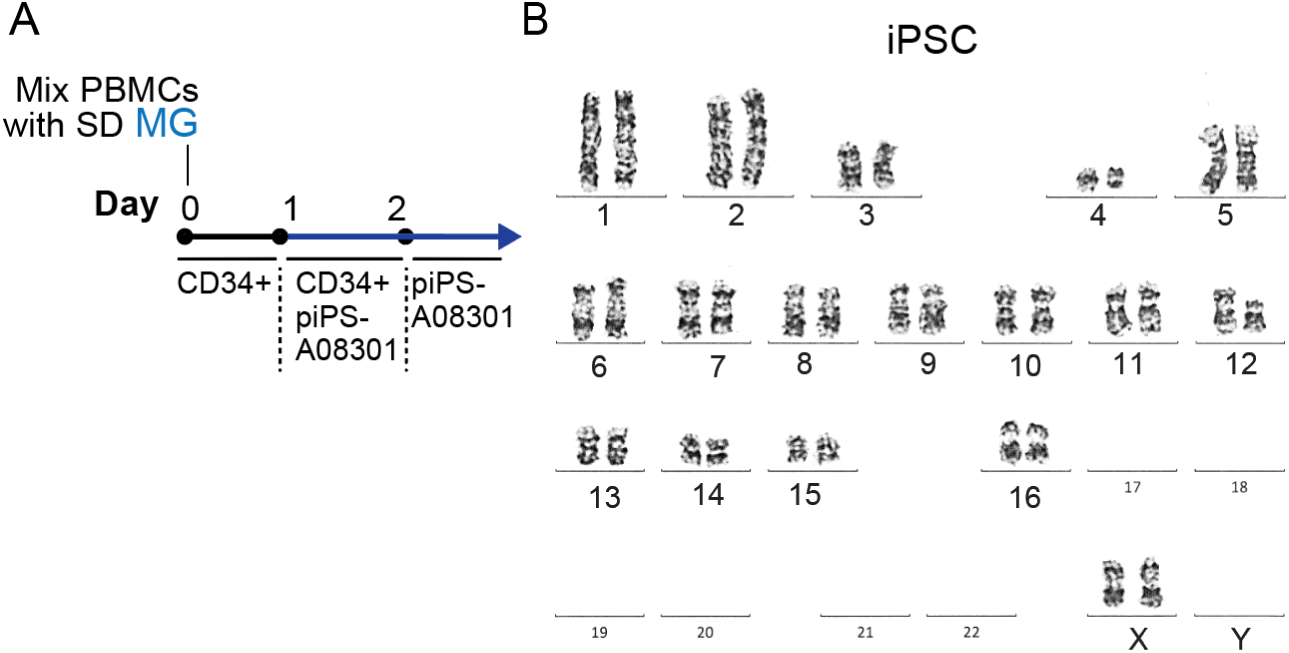
Reprogramming strategy and characterization of Red River Hog (Potamochoerus porcus) iPSCs. (A) Schematic timeline of the Sendai virus-mediated reprogramming protocol for Red River Hog peripheral blood cells. (B) Karyotype analysis of iPSC^Rrh^ derived from blood cells, confirming a normal female karyotype (34, XX).

**Supplementary Figure 5.**
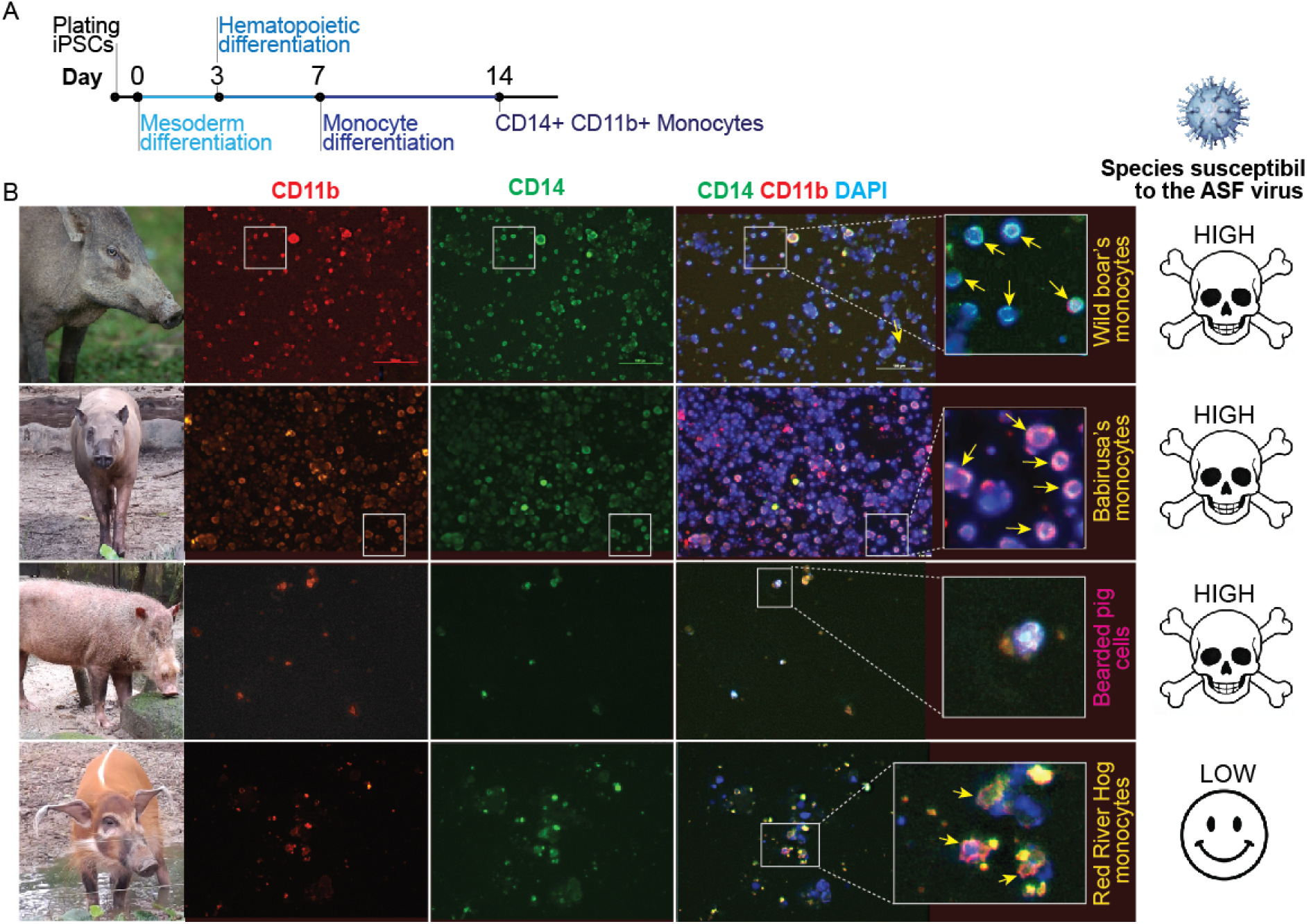
Differentiation of porcine iPSCs into CD14⁺CD11b⁺ monocytes. (A) Schematic timeline of the monocyte differentiation protocol. (B) Immunofluorescence staining for the monocyte surface markers CD14 and CD11b in cells harvested every 2–3 days between days 17 and 27 of differentiation, shown for iPSC^W^ (wild boar; high ASF susceptibility), iPSC^Ba^ (Babirusa; high ASF susceptibility), iPSC^Bp^ (Bornean bearded pig; high ASF susceptibility), and iPSC^Rrh^ (Red River Hog; low ASF susceptibility). Scale bars: 100 μm.

## Notes

### Competing Interest Statement

The authors have declared no competing interest.

### Summary of Updates

Add an author - Nicole Liling TAY

